# EVRC: Reconstruction of chromosome 3D structure models using Error-Vector Resultant algorithm with Clustering coefficient

**DOI:** 10.1101/2023.05.11.540436

**Authors:** Xiao Wang, Jie Li, Wei-Cheng Gu, Bin-Guang Ma

## Abstract

Reconstruction of 3D structure models is of great importance for the study of chromosome function. In this paper, we present a novel reconstruction algorithm, called EVRC, which utilizes co-clustering coefficients and error-vector resultant for chromosome 3D structure reconstruction. To evaluate the effectiveness and accuracy of the EVRC algorithm, we applied it to simulation datasets and real human Hi-C datasets. The results show that the reconstructed structures have high similarity to the original/real structures, indicating the effectiveness and robustness of the EVRC algorithm. Furthermore, we applied the algorithm to the 3D conformation reconstruction of the wild-type and mutant *Arabidopsis thaliana* chromosomes and demonstrated the differences in structural characteristics between different chromosomes. We also accurately showed the conformational change in the centromere region of the mutant compared with the wild-type of *Arabidopsis* chromosome 1. Our EVRC algorithm is a valuable software tool for the field of chromatin structure reconstruction, and holds great promise for advancing our understanding on the chromosome functions.

## 1. Introduction

Since the implementation of ENCODE, scientists have discovered that the folding and winding of chromatin in the nucleus is not random. Regulatory elements require physical contact with target genes to perform their functions [1-3]. The high-level structure of chromatin is increasingly recognized as a crucial component of many nuclear activities [3, 4]. While traditional fluorescence microscopic imaging techniques can directly observe the position and relative distance of different chromosomes in the cell, they suffer from low flux and low resolution. However, the birth of chromosome conformation capture (3C) technology and its derivative technologies has provided researchers with a way to obtain a large number of chromatin interaction data [5, 6]. Researchers have discovered that the 3D structure of chromatin plays an important role in many life activities, such as gene expression regulation [2, 7], cell development [8], and the occurrence of genetic diseases [9, 10]. By using chromatin interaction data to reconstruct chromosome 3D structure models, researchers can gain a deeper understanding of the fine structure and dynamic changes of chromosomes. This will enable researchers to explore the chromatin formation mechanism of the genome, discover the relationship between regulatory function and spatial distribution, and provide new ideas for the study of heterosis, genetic diseases, and cancer cell therapy.

The current methods for modeling the 3D structure of chromatin can be broadly categorized into two types: thermodynamics-based (TB) and restraint-based (RB) modeling [11]. TB modeling uses polymer physical properties of chromatin fibers to construct the spatial arrangement of chromatin, while RB modeling employs spatial constraints between chromatin fragments obtained by Hi-C technology to build 3D models of chromatin [12]. One of the advancements in this field is the miniMDS method, which uses genome partitioning and parallelization to reduce memory requirements and achieve faster speed. This technique can calculate the 3D structure of the human genome at kilobase resolution in less than five hours [13]. ClusterTAD is another technique that uses unsupervised machine learning models and image analysis technology to identify TADs (topologically associating domains) in the diagonal of the interaction heat map; the interaction matrix is treated as pixel points in the image to identify the TADs [14]. HiCExplorer, on the other hand, utilizes machine learning methods to identify TAD boundaries accurately; it distinguishes TAD boundaries from non-boundaries and detects the ones missed when using Hi-C data alone [15]. Researchers have also incorporated information obtained from other experimental methods in structure modeling, such as FISH experiments [16], which detect relative positions and spatial distances between different DNA fragments. These integrations can help improve and evaluate the reliability and stability of chromosome 3D structure reconstruction. Recently, more and more reconstruction tools have be developed [17-19]. For example, ShNeigh combines the classical MDS (multidimensional scaling) technique with local dependence of neighboring loci to infer the 3D structure from noisy and incomplete contact frequency matrices [17].

In this paper, we present a novel EVRC algorithm that utilizes Hi-C experiments data to reconstruct the 3D structure of chromatin. Our approach relies on the co-clustering coefficient and error-vector resultant [20]. Specifically, the algorithm begins by calculating the co-clustering coefficient between chromatin fragments. It then adds the error vector of each fragment weighted by the co-clustering coefficient together to iteratively optimize the reconstruction of the 3D chromatin structure. We applied the EVRC algorithm to six simulated structures and real Hi-C data sets to demonstrate its validity and accuracy. Our results indicate that the algorithm is an effective tool for reconstructing the chromatin structure and has the potential to significantly advance the study of chromatin structure and function.

## 2. Materials and method

### 2.1 Data source

#### 2.1.1 Simulation data

To assess the effectiveness of EVRC algorithm for 3D structure reconstruction, we generated multiple simulation datasets with six 3D structure models of varying complexities: (A) circular curve (circle), (B) open spiral curve (spiral), (C) closed spiral curve (circular spiral), (D) replication fork, (E) double helix, (F) double spherical helix. (A), (B), (C) are single-chain structures, and (D), (E), (F) are double-chain structures, simulating different chromatin morphology of prokaryotes/eukaryotes with one or multiple chromosomes. For the double-chain model structures, the intra-chain and inter-chain contacts should be considered, which simulates the intra- and inter-DNA interactions in chromosomes. Each structure is composed of 500 points. In the 3D structure, the reciprocal of the space distance between each two points is taken as the interaction frequency between the two points, and the interaction matrix of the structure is obtained in this way. The single-chain structure generates only the interaction matrix within a single curve (simulating single chromosome), while the double-chain structure generates the interaction matrix within each of the two curves and the interaction matrix between them (simulating multiple chromosomes).

#### 2.1.2 Hi-C data

We used the published chromosome interaction data of two species: the raw chromosome interaction frequency matrix with 50kb resolution of human IMR90 cell line (GEO accession number: GSE63525)[6], and normalized chromosome interaction frequency matrix with 50kb and 200kb resolution of *Arabidopsis* (GEO accession number: GSE37644, where GSM1078404 was the wild-type Hi-C data and GSM1078405 was the AtMORC6 mutant Hi-C data) [21].

#### 2.1.3 FISH data

The FISH data for human cell line IMR90 used in this study was adopted from a previous work [16], which is originally from https://www.sciencemag.org/content/353/6299/598/suppl/DC1 [22].

### 2.2 The EVRC algorithm

#### 2.2.1 Conversion of interaction frequency to spatial distance

The Hi-C experimental data generates a chromatin interaction matrix that reflects the intensity of interactions between DNA fragments (bins). The closer the space distance between bins, the higher the interaction intensity. In order to transform the interaction intensity into spatial distance information, the commonly used negative exponential function is adopted in this paper:

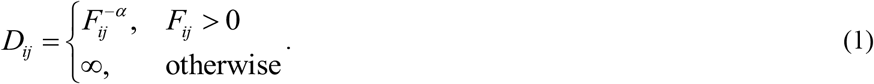

Here, *D*_*ij*_ represents the spatial distance between bin_*i* and bin_*j, F*_*ij*_ represents the interaction frequency between bin_*i*_ and bin_*j*_, and *α* is the exponential parameter. In this paper, the *α* value defaults to 0.5 and can be adjusted according to the data characteristics.

#### 2.2.2 Co-clustering coefficient

In graph theory, the clustering coefficient (*CC*) is used to describe the degree of clustering between nodes in a graph [23]. Specifically, it measures the closeness of the connections between the adjacent nodes of a node. In an undirected network, if node *i* has *k*_*i*_ neighbor nodes, and if all the *k*_*i*_ nodes are connected in pairs, the total number of edges *E*_*i*_ would be: *E*_*i*_ = *k*_*i*_(*k*_*i*_ - 1)/2. If the actual number of edges between the *k*_*i*_ nodes is *e*_*i*_, then the clustering coefficient of node *i* is:

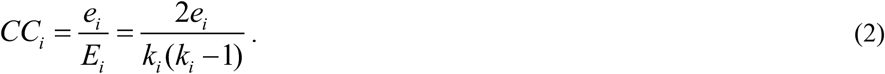

From this formula, it can be seen that the higher the number of edges actually present between the neighbor nodes, the greater the clustering coefficient of node *i* (0 ≤ *CC*_*i*_ ≤ 1), indicating a closer group.

The chromatin interaction matrix reflects the interaction relationships between the DNA fragments (bins) and can be regarded as an undirected network among bins, in which each bin is a node. After conversion to a distance matrix, *D*_*ij*_ represents the spatial distance between bin_*i*_ and bin_*j*_. If *D*_*ij*_ is not ∞, there is a connection between bin_*i*_ and bin_*j*_. Based on Eq. (2), we can define the co-clustering coefficient between bin_*i*_ and bin_*j*_. Let the set of neighbor nodes of node bin_*i*_ be N_*i*_, and the set of neighbor nodes of node bin_*j*_ be N_*j*_. The union of the neighbor sets of node bin_*i*_ and node bin_*j*_ is U_*ij*_ = N_*i*_ ∪ N_*j*_, which contains *k*_*ij*_ nodes. The co-clustering coefficient of the nodes bin_*i*_ and bin_*j*_ is defined as:

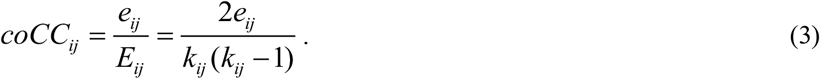

In this formula, *coCC*_*ij*_ represents the co-clustering coefficient of nodes bin_*i*_ and bin_*j*_ (**Figure 1**); *e*_*ij*_ is the actual number of edges connecting the nodes in U_*ij*_, and *E*_*ij*_ is the number of edges if all nodes in U_*ij*_ are connected in pairs. The closer the *coCC*_*ij*_ to 1, the greater the tendency of co-clustering between bin_*i*_ and bin_*j*_ in the 3D structure of chromatin. For example, bins in the same TAD tend to have higher co-clustering coefficients. We consider the co-clustering coefficient as an important parameter for 3D structure reconstruction.

**Figure 1.**
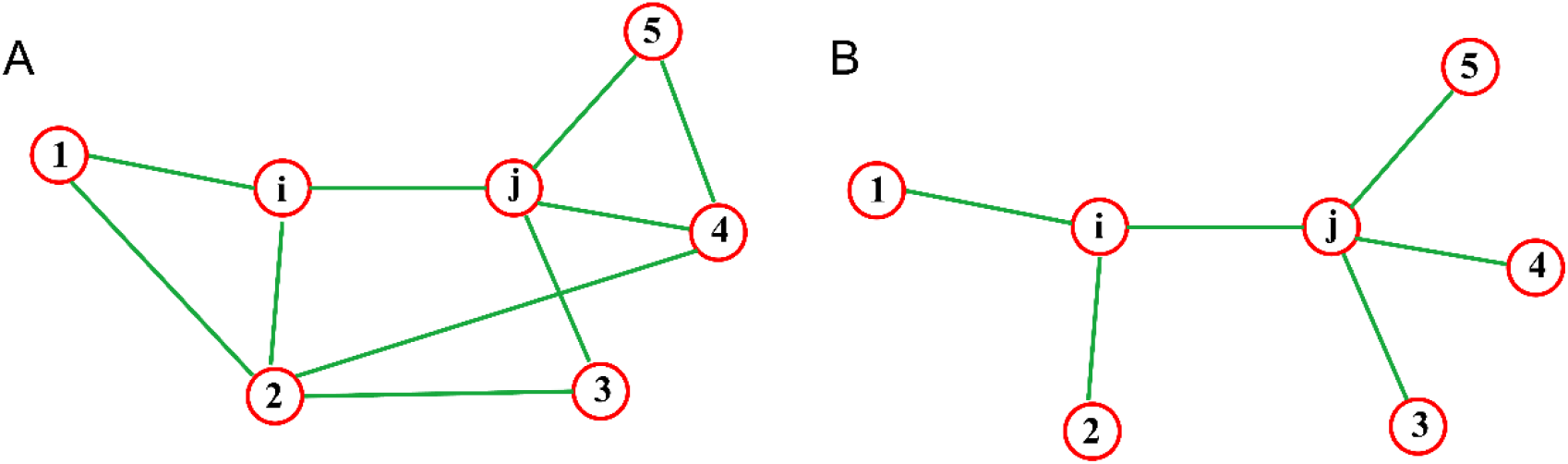
Definition of co-clustering coefficient. Red circles represent nodes, and green lines represent interactions between nodes. In the subplot A, there are more connecting edges between the neighbor nodes of node *i* and node *j*, so the co-clustering coefficient is large (*coCC*_*ij*_ = 10/21 = 0.476). In the subplot B, the number of connecting edges between the neighbor nodes of nodes *i* and *j* is small, so the co-clustering coefficient is small (*coCC*_*ij*_ = 6/21 = 0.286).

#### 2.2.3 Reconstruction of 3D chromosome structure model

The objective of restraint-based chromosome 3D structure modeling algorithm is to generate a 3D structure that fits experimental data as closely as possible. In previous work, we proposed an algorithm based on the Error Vector Resultant (EVR) to model the 3D structure of chromosomes [20]. In this paper, we have developed the EVR algorithm by incorporating the co-clustering coefficient between bins, resulting in the EVRC algorithm. For each bin, its initial 3D coordinates are randomly generated. Assuming the position relationship between bin_*i*_ and bin_*j*_ (*i* ≠ *j*) as shown in **Figure 2A**, *P*_*i*_ and *P*_*j*_ represent the corresponding position vectors, *e*_*ij*_ represents the unit vector of *p*_*i*_ - *p*_*j*_, and *D*_*ij*_ represents the spatial distance between bin_*i*_ and bin_*j*_ (*D*_*ij*_ ≠ ∞). The error vector *E*_*ij*_ between bin_*i*_ and bin_*j*_ is defined as:

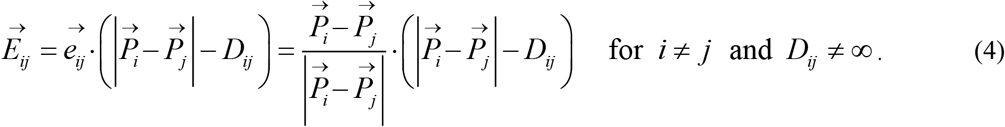

**Figure 2.**
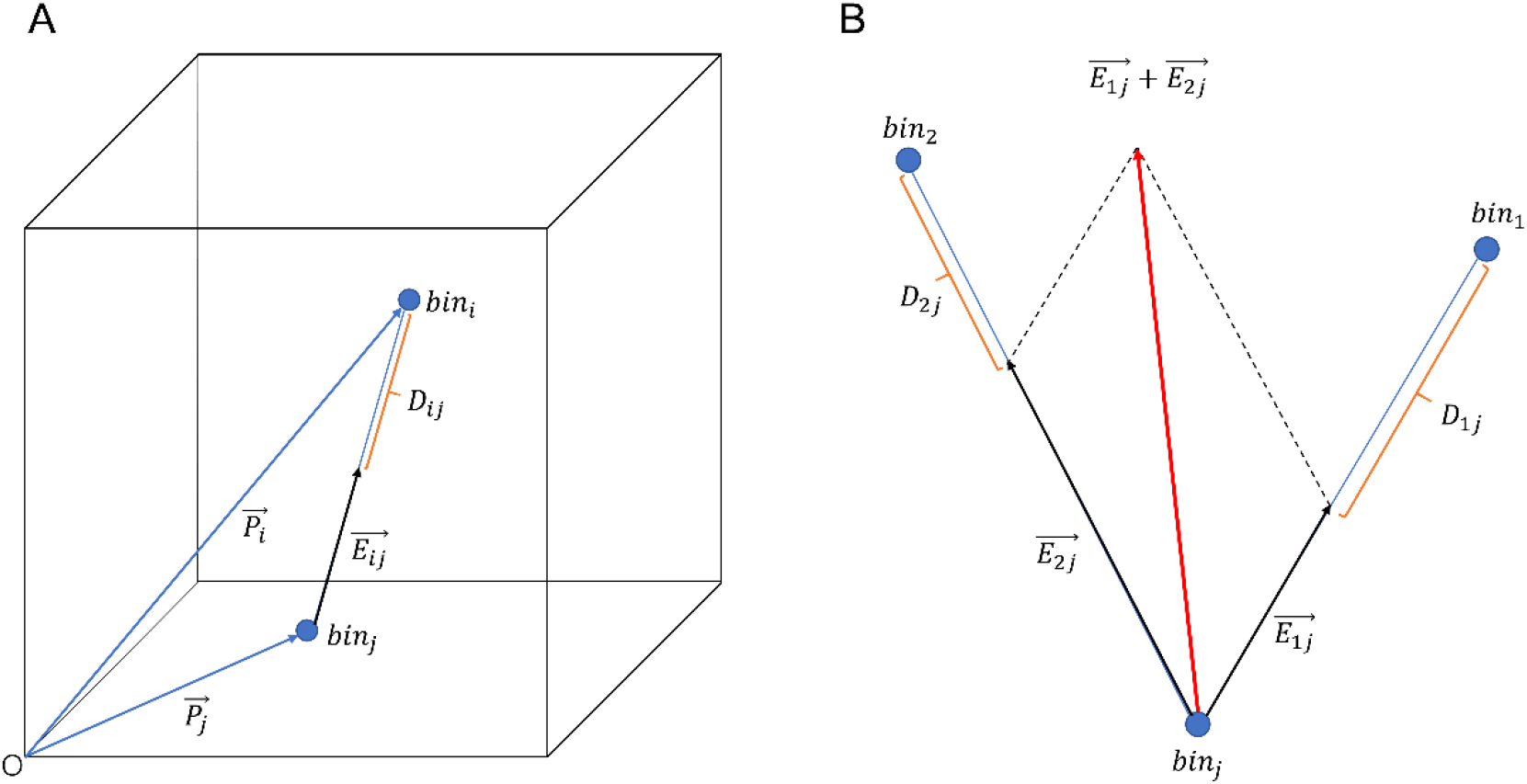
Diagram of error-vector resultant. (A) The error-vector *E*_*ij*_ between bin_*i*_ and bin_*j*_. (B) The resultant of error-vectors of *E*_1*j*_ and *E*_2*j*_ according to the parallelogram rule.

The sum of the error vectors from bin_*j*_ to other bins can be obtained using the parallelogram rule. For example, the sum of the error vectors from bin_*j*_ to bin_1_ and bin_2_ is shown in **Figure 2B**. Summing over the genome-wide range, the total error vector of bin_*j*_ is:

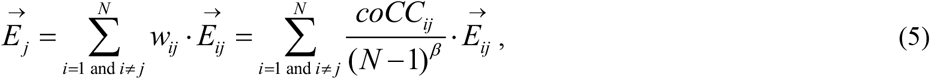

where *w*_*ij*_ = *coCC*_*ij*_ / (*N* - 1)^*β*^ is the weight in sum, *coCC*_*ij*_ is the co-clustering coefficient between bin_*i*_ and bin_*j*_, *N* is the total number of bins, and *β* is a parameter for adjusting the convergence speed (called convergence factor; the larger its value, the slower the convergence speed) that is set to 0.1 by default.

According to the flow of EVR algorithm [20], the structure model of chromosome is obtained through an iterative process. In each iteration step, bin_*j*_’s resultant error vector *E*_*j*_ is added to its current 3D coordinates to obtain the new coordinates and then the iteration proceeds to the next step. For all bins in the genome, the global optimization objective is defined as:

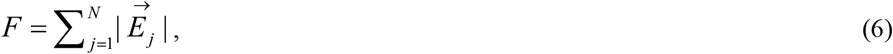

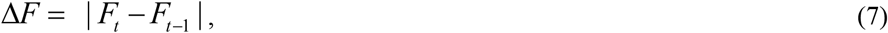

where, *F* is the sum of resultant error vectors of all bins, and *F*_*t*_ and *F*_*t*-1_ are the *F* values of two successive iterative steps *t* and *t*-1. With continuous iterative optimization, the resultant error vector of each bin and the *F* value become smaller and smaller, making the generated 3D structure more and more consistent with the interaction data. According to the flow of EVR algorithm [20], when Δ*F* is less than a certain value or the maximum number of iteration steps is reached, the iterative process stops, resulting in the final chromosome 3D structure model. To better visualize the structure model, the 3D coordinates of the structure can be smoothed using a linear Gaussian filtering function, which contains a parameter (smoothing factor) that is set to ‘auto’ by default, meaning no smoothing. Users can adjust the value of this parameter according to their needs. Gaussian smoothing is only used to optimize visualization, not for algorithm evaluation, that is, algorithm evaluation is always based on unsmoothed coordinates.

### 2.3 Model evaluation indexes

To assess the accuracy of the reconstruction algorithm, simulation data generated based on model structures are often employed. In this paper, the 3D structures are reconstructed from the simulation data of six models mentioned above and then compared with the original model structures to evaluate the effectiveness of the reconstruction algorithm. Two indicators are used to measure the similarity between the reconstructed 3D structure and the original structure: Root Mean Squared Deviation (RMSD) of 3D coordinates [24] and Pearson Correlation Coefficient (PCC) based on spatial distance [20].

## 3. Results

### 3.1 Algorithm demonstration

#### 3.1.1 Iteration and convergence

The EVRC algorithm initializes the chromosome structure by randomly assigning 3D coordinates to each bin. Then, under the guidance of the resultant error vector, the 3D structure is gradually optimized through an iterative process until it converges or reaches the maximum number of iterations. In each iteration step, the values of *F* and Δ*F* are calculated according to equations (6) and (7), respectively. Under default parameter settings, the curve of *F* value changing with the number of iterations for the six simulating structures is shown in **Figure 3**. As can be seen, the *F* value of the randomly generated initial structure is high, but decreases rapidly after several iterative steps, and tends to zero steadily after some intermediate fluctuations. The iterative process terminates when the convergence criterion (Δ*F* < 1E-6) is met, and the optimized structure is obtained (**Figure 4**). The reconstructed 3D structure is highly consistent with the original structure, and the RMSD values are small. The results demonstrate that the EVRC algorithm is effective for reconstructing both open-loop and closed-loop structures, as well as single-chain and double-chain structures. Therefore, the EVRC algorithm is generally effective for 3D chromatin reconstruction.

**Figure 3.**
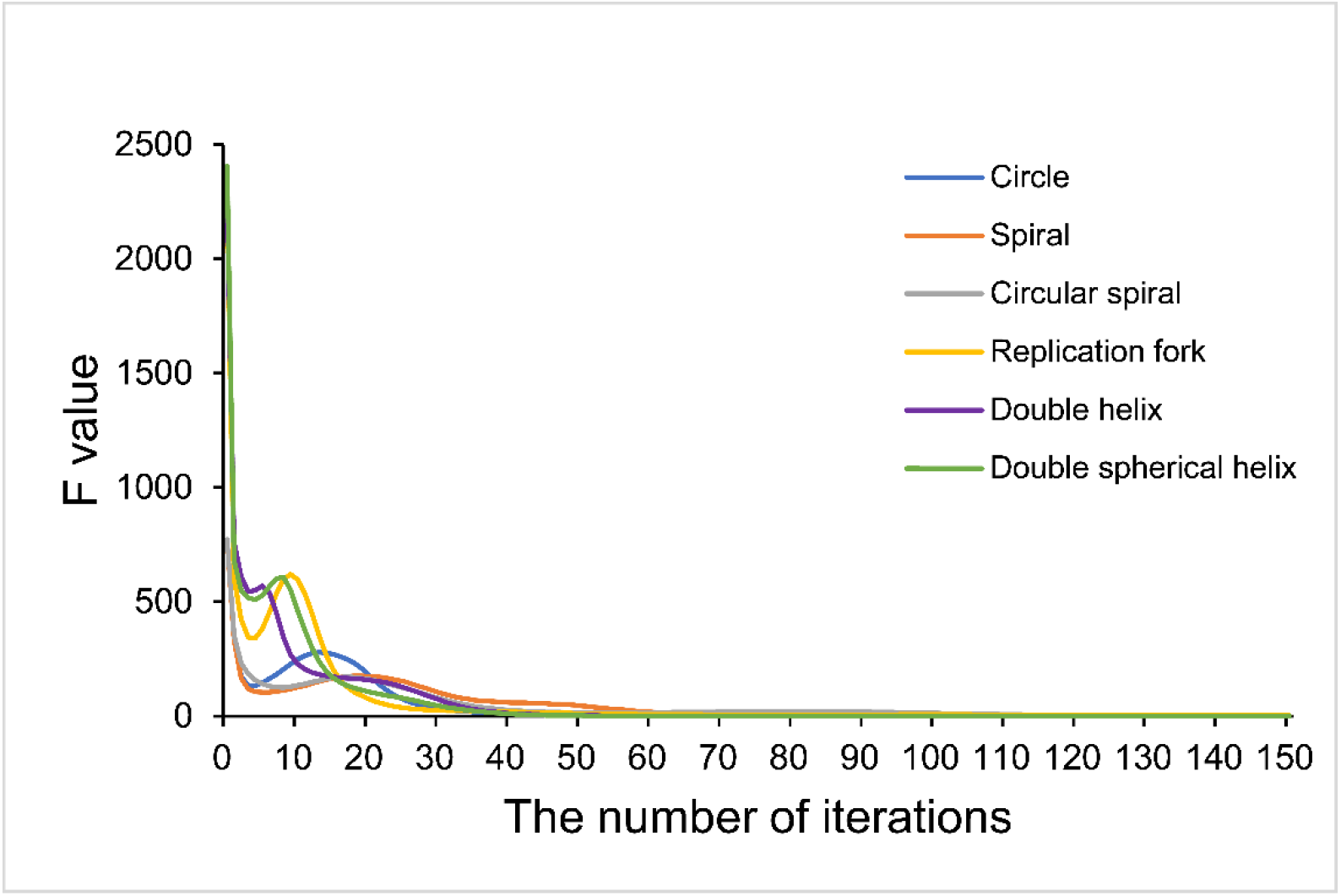
Changes of *F* value with the number of iterations in the process of iterative optimization.

**Figure 4.**
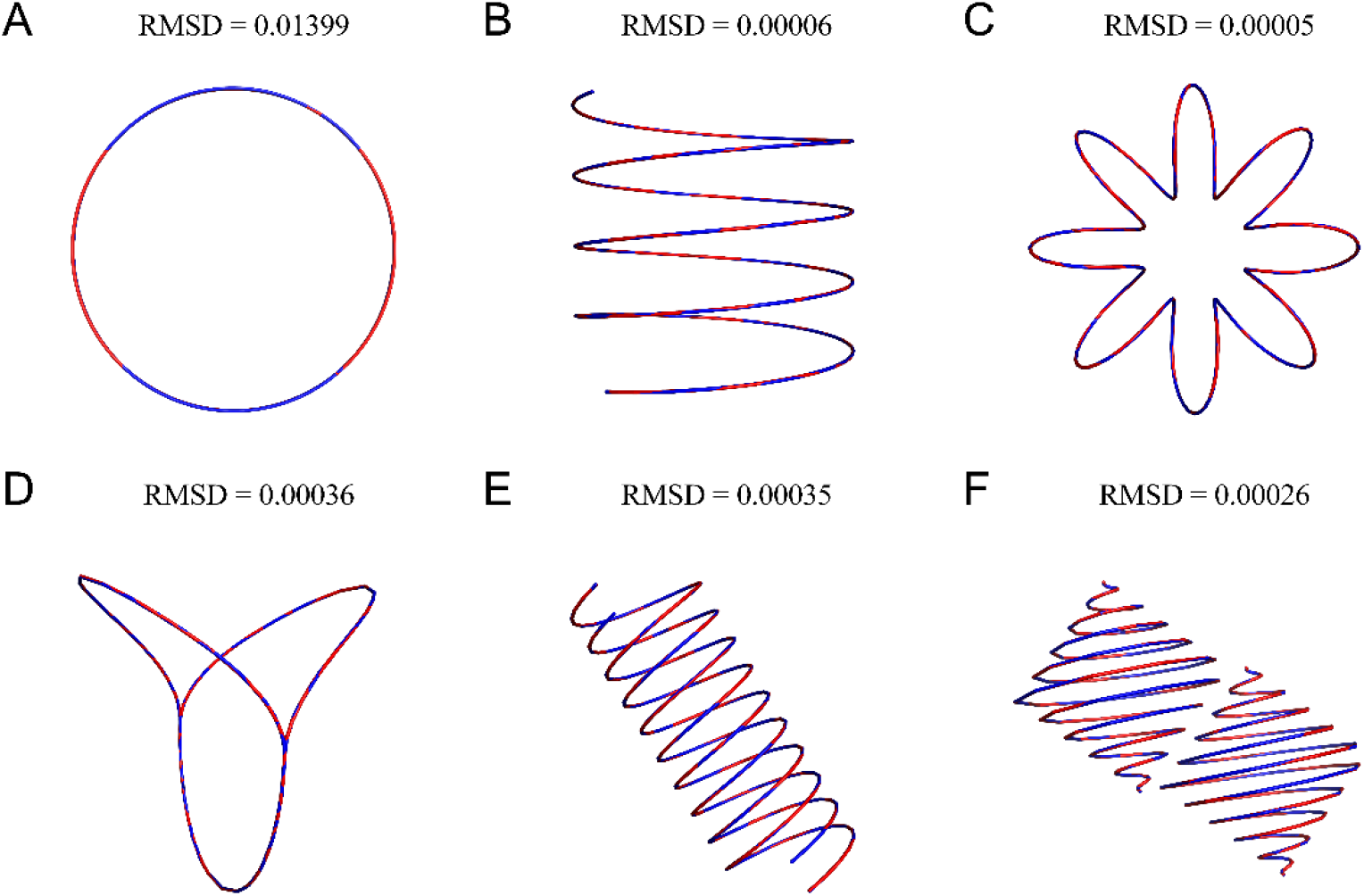
The reconstructed structure (blue curve) is highly consistent with the real structure (red curve). (A) circular curve (circle); (B) open spiral curve (spiral); (C) closed spiral curve (circular spiral); (D) replication fork; (E) double helix; (F) double spherical helix.

#### 3.1.2 Noise and smoothing

To demonstrate the effectiveness of EVRC algorithm for reconstructing 3D structures under different levels of noise, we added noise to the simulation data and reconstructed the six simulating structures based on the noisy data. **Figure 5** displays the changes in RMSD and PCC with increasing noise level. The figure shows that the RMSD value gradually increases with increasing noise level, while the PCC decreases. At a noise level of 0.1, the PCCs of all the six simulating structures are greater than 0.99, indicating the high similarity with the real structures. At the maximum noise level of 1, the corresponding PCCs are still all greater than 0.96. The reconstructed 3D structure at noise level 1 (**Figure 6**) shows that although the curves are fluctuated due to the high noise, they still closely surround the real structure without significant deviation. Our modeling scheme assumes that each bin corresponds to a point in the 3D structure of a chromosome, which results in a relatively sharp curve formed by directly connecting the points with straight lines. For a better visualization of the reconstructed structure, Gaussian filtering function might be used for smoothing as shown in **Figure S1**, where the noise level is 1 and the smoothing factor is 2.

**Figure 5.**
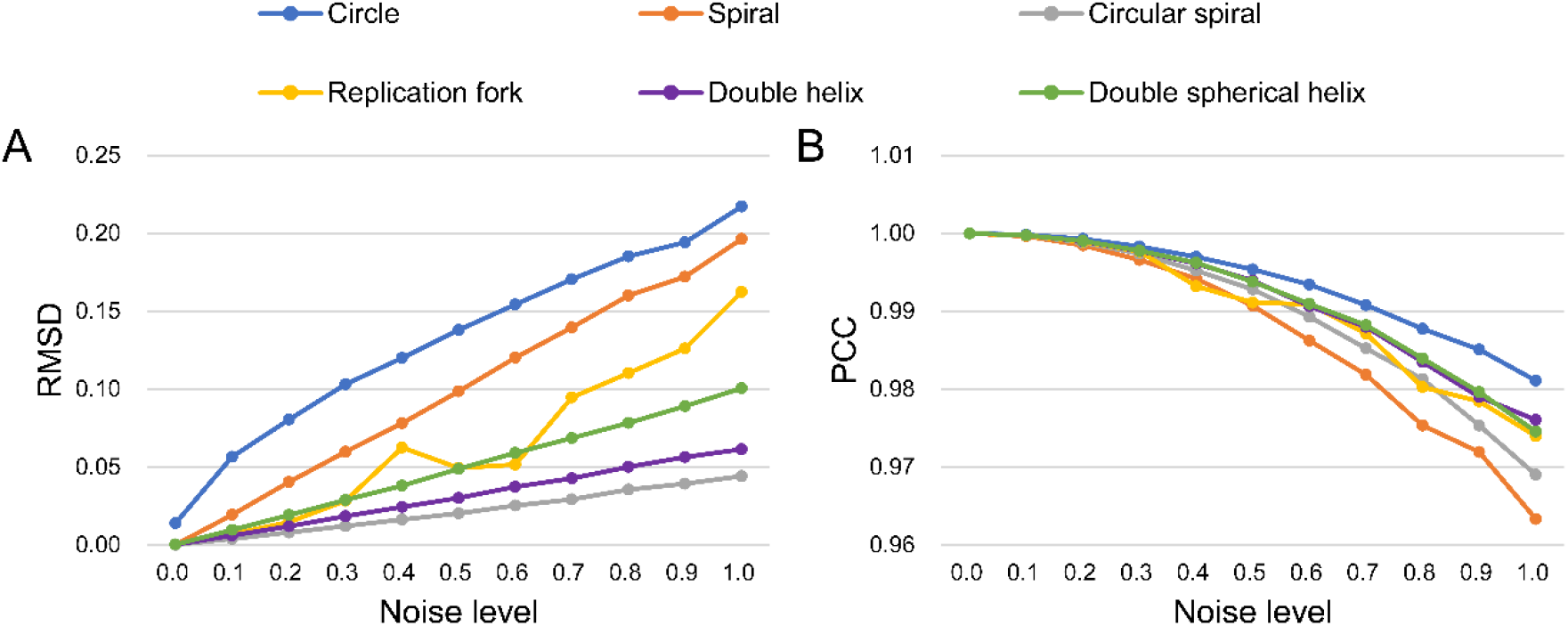
Performance of EVRC algorithm reconstruction of the six simulating structures under different noise levels. (A) The variation of RMSD with noise level; (B) The variation of PCC with noise level.

**Figure 6.**
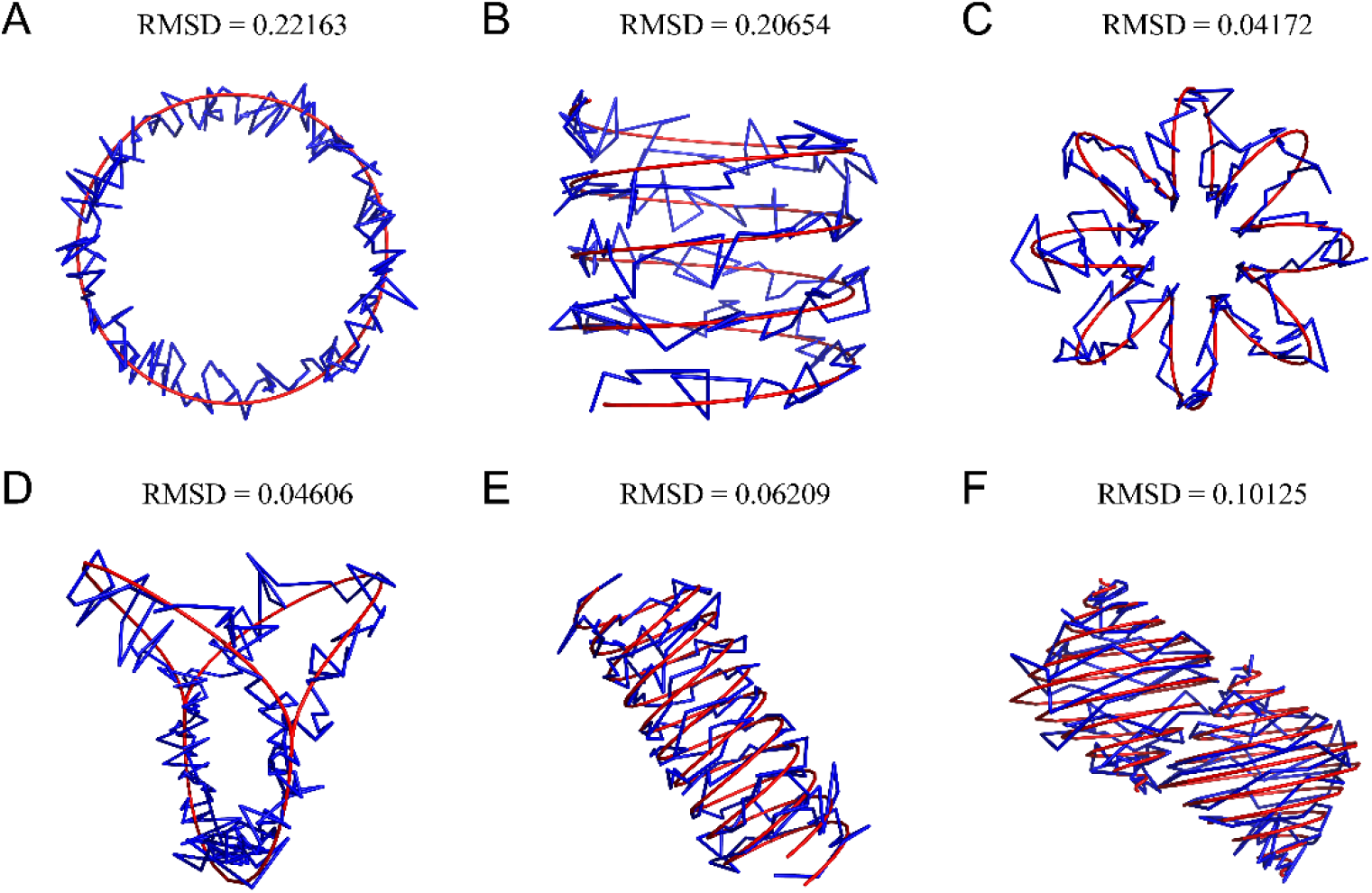
Comparison between reconstructed structure (blue curve) and original structure (red curve) when noise level is 1. (A) circular curve (circle); (B) open spiral curve (spiral); (C) closed spiral curve (circular spiral); (D) replication fork; (E) double helix; (F) double spherical helix.

### 3.2 Algorithm evaluation

#### 3.2.1 Simulation data

We compared the performance of EVRC with four representative algorithms, namely miniMDS [13], ShRec3D [25], ShNeigh [26] and MOGEN [27] on the six simulating structures with varying noise levels. All the algorithms were run with default parameters. To reduce the impact of randomness, each algorithm was run ten times at each noise level to obtain the average RMSD and PCC. The results indicate that when the noise level is near 0, the RMSD values of EVRC, ShRec3D and ShNeigh are also close to 0 (**Figure 7**), while the PCCs of these algorithms are close to 1 (**Figure 8**) for the single-chain structures. This indicates that these algorithms can effectively reconstruct the original structure at very low noise levels. However, as the noise level increases, the RMSD values of most algorithms increase gradually, while the PCC values decrease. Notably, EVRC displays better stability than other algorithms. Thus, the EVRC algorithm exhibits superior stability compared to other algorithms as the noise level increases.

**Figure 7.**
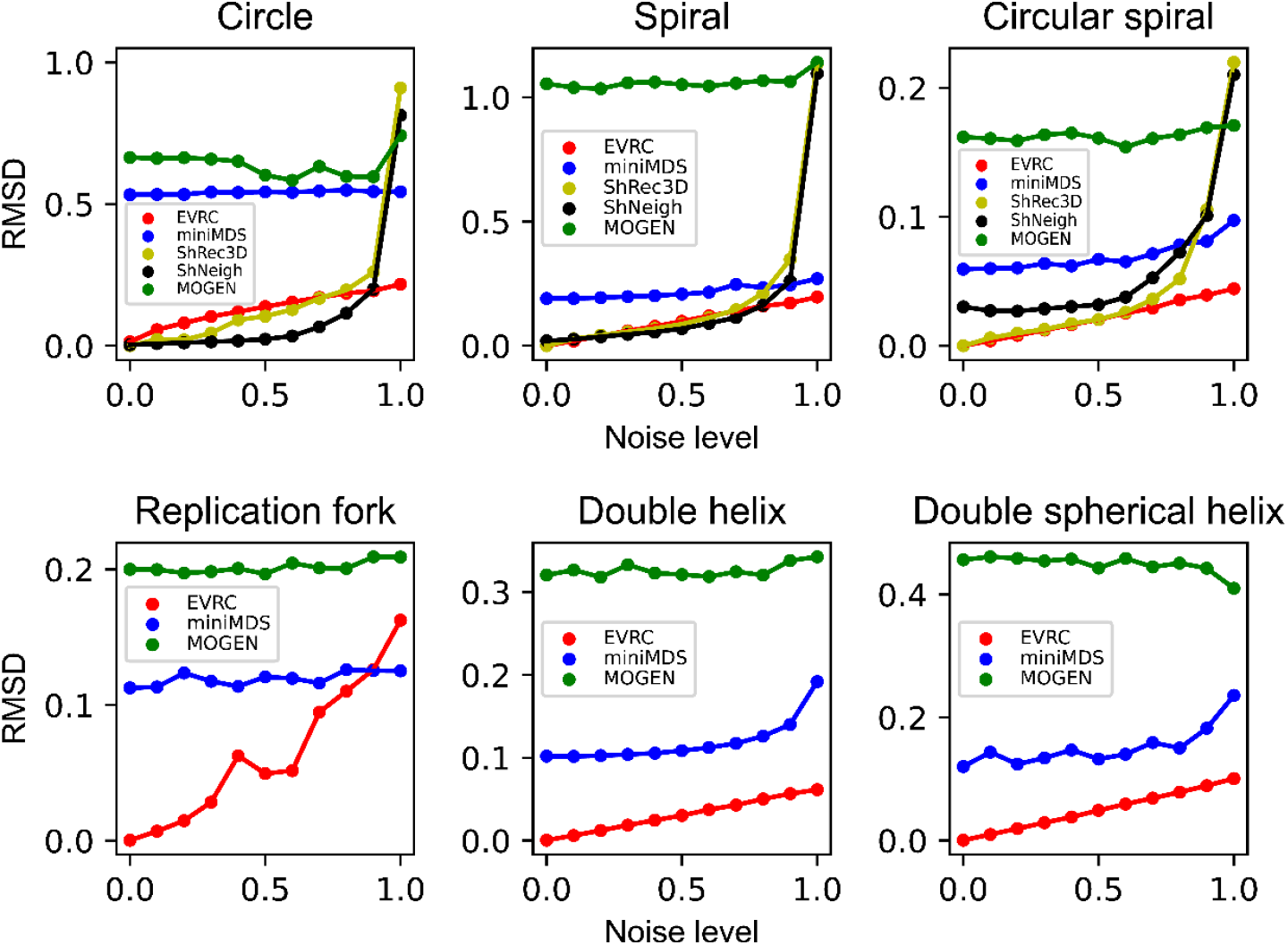
Based on the noisy data of six simulating structures, the RMSD values of the reconstructed structures by each algorithm compared with the original structures under different noise levels. Note that only EVRC, miniMDS and MOGEN have reconstruction results for the double-chain structures (namely, replication fork, double helix, and double spherical helix).

**Figure 8.**
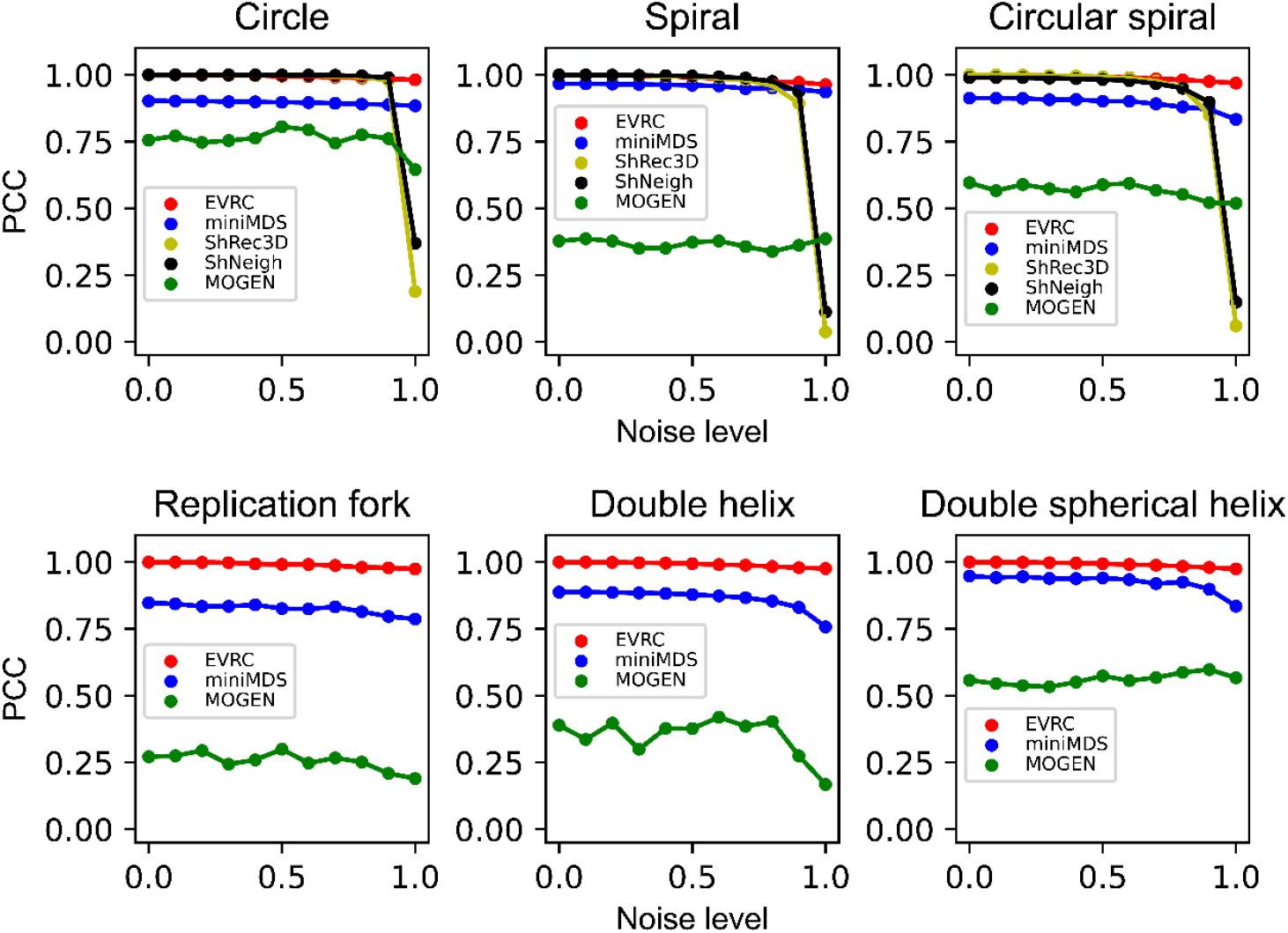
Based on the noisy data of six simulating structures, the PCC values between the spatial distances in the reconstructed structures and in the original structures under different noise levels. Note that only EVRC, miniMDS and MOGEN have reconstruction results for the double-chain structures (namely, replication fork, double helix, and double spherical helix).

#### 3.2.2 Real Hi-C data

In addition to the simulating structures, we also evaluated the performance of each algorithm using datasets from the real Hi-C experiments of human cell lines. Given that the real structure of human chromosomes is unknown, we cannot calculate the RMSD value between the reconstructed structure and the real structure of human chromosomes. However, we can use the FISH experimental data to evaluate the accuracy of the reconstruction algorithm. Specifically, we found the Hi-C data [6] and the corresponding FISH data [22] on chromosomes No. 20 and No. 21 of the human IMR90 cell line through a literature search. We mapped the fluorescent-labeled sites onto the reconstructed 3D structure, calculated the spatial distances between these sites in the reconstructed structure, and analyzed the correlation with the distances measured in the FISH experiments. The results, as shown in **Table 1**, revealed that, the PCC of EVRC is 0.7576 for chromosome No. 20 and 0.8206 for chromosome No. 21, the highest among the compared algorithms. Meanwhile, for chromosome No. 20, the miniMDS and ShRec3D methods failed to generate 3D structure due to the missing interaction data for the middle region of this chromosome in the Hi-C experiment. Overall, these results demonstrate that the EVRC algorithm is accurate.

**Table 1.** Pearson correlation coefficient between the spatial distance in reconstructed structure and the distance measured by FISH experiment in human IMR90 cells.

| Chromosome number | Algorithm | PCC | *P*-value |
| --- | --- | --- | --- |
| 20 | EVRC | 0.7576 | 3.08E-82 |
|  | ShNeigh | 0.6861 | 8.42E-62 |
|  | MOGEN | 0.2378 | 5.20E-07 |
| 21 | EVRC | 0.8206 | 6.46E-130 |
|  | ShNeigh | 0.8140 | 3.56E-126 |
|  | ShRec3D | 0.8187 | 8.22E-129 |
|  | miniMDS | 0.7989 | 3.22E-118 |
|  | MOGEN | 0.3618 | 9.01E-18 |

### 3.3 Application of EVRC to plant 3D genomes

Chromosome conformation capture technologies have led to the publication of chromatin interaction data for more and more species. In this study, we applied the EVRC algorithm to the chromatin interaction datasets of *Arabidopsis* to reconstruct the 3D chromosome structures. The *Arabidopsis thaliana* chromatin interaction dataset used in this study was derived from two different samples, GSM1078404 (wild-type) and GSM1078405 (AtMORC6 mutant), both available in the GEO database (accession number: GSE37644). The datasets included five intra- and inter-chromosome interaction matrices. AtMORC6 is a member of the conservative Microrchidia (MRC) family that catalyzes changes in chromosome superstructure [21]. AtMORC6 mutants in *Arabidopsis thaliana* show depolymerization of heterochromatin in centromeres, leading to an increased interaction between the centromeric region and the rest of the genome [21].

The input data for the EVRC algorithm was the normalized interaction matrixes with a resolution of 50 kb and 200 kb. The 3D structure of chromatin was obtained by running 20,000 iteration steps under default parameters. **Figure 9** shows the reconstructed 3D conformation of chromosomes for wild-type *Arabidopsis thaliana*. The colors green, cyan, magenta, yellow and brown represent chromosomes 1 to 5, respectively. Figure 9A has a resolution of 200 kb, and Figure 9B has a resolution of 50 kb. The overall morphology of the chromosomes is consistent between different resolutions, with the structure in Figure 9A being smoother and the structure in Figure 9B being more detailed. These results indicate that the EVRC algorithm accurately reconstructs chromosome conformations at different resolutions with strong stability. The shapes of the five chromosomes are different, with chromosomes 1, 3, and 5 showing a narrow “U” shape, and the centromere region having varying degrees of curvature. Chromosomes 2 and 4 are hooked, with the ends of the chromosomes being closer. The common feature for all the five chromosomes is the loose centromere region. The 3D reconstruction results of the five chromosomes in *Arabidopsis thaliana* demonstrate the effectiveness of the EVRC algorithm in reconstructing conformation of multiple chromosomes, suggesting that EVRC can serve as a valuable chromatin 3D reconstruction tool in eukaryotes.

**Figure 9.**
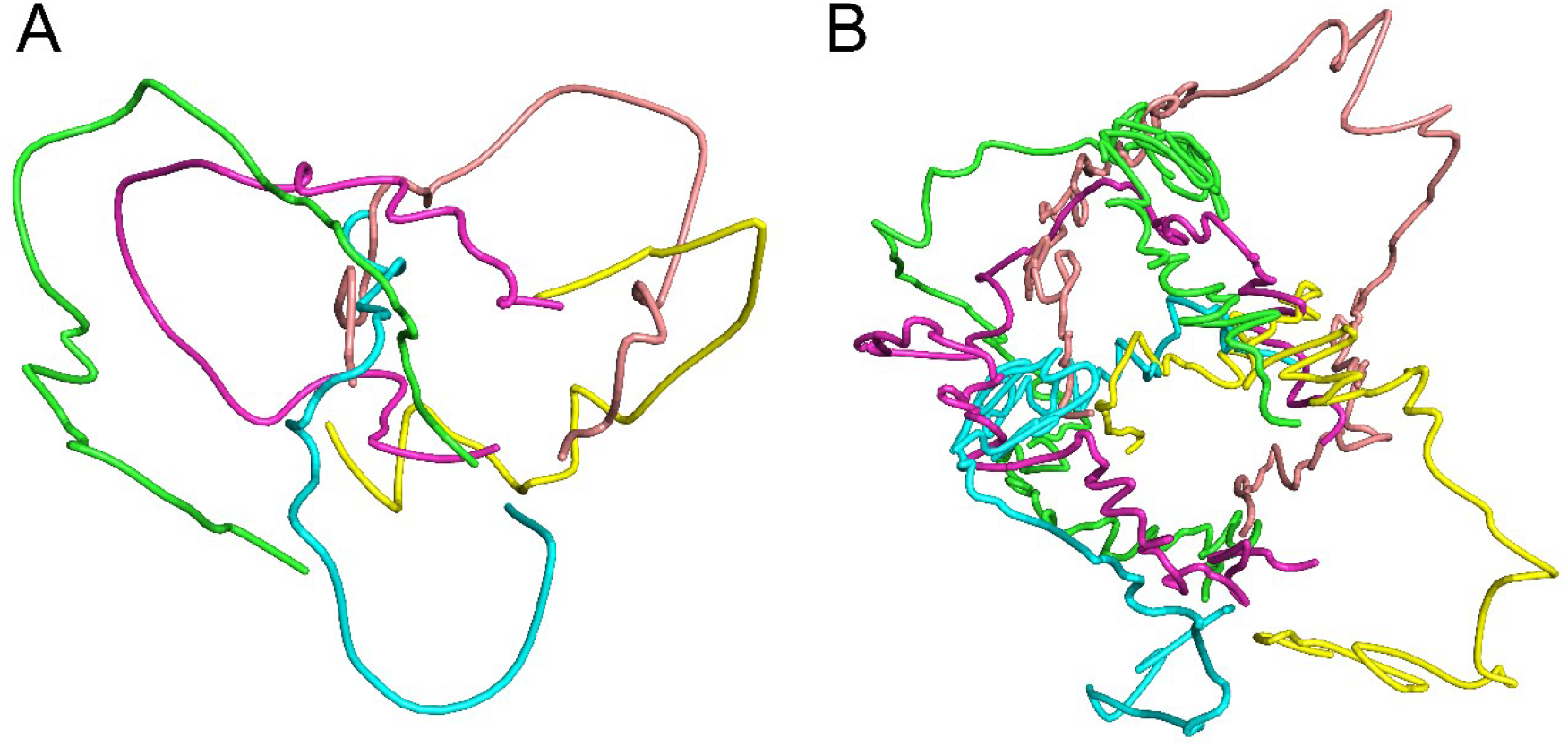
The 3D conformation of wild-type *Arabidopsis thaliana* chromosomes reconstructed by EVRC algorithm. Chromosomes 1 to 5 are shown in green, cyan, magenta, yellow and brown, respectively. (A) at a resolution of 200 kb; (B) at a resolution of 50 kb.

Figure 10. depicts the 3D conformation of chromosome 1 at a resolution of 50 kb, with the wild-type shown in red and the mutant in blue. The non-overlapping part represents the centromere region. As seen in the figure, the structures of the wild-type and the mutant show a high degree of similarity except for the centromere region, which is separated by 180°. In the mutant, the centromere region is closer to the surrounding chromatin, resulting in enhanced interaction and giving the chromosome a “C” shape. In the wild-type, the centromere region folds upward, giving the chromosome an “S” shape. The reconstruction results are consistent with the functional characteristics [21], indicating the validity and accuracy of the EVRC algorithm in plant chromosome structure reconstruction.

**Figure 10.**
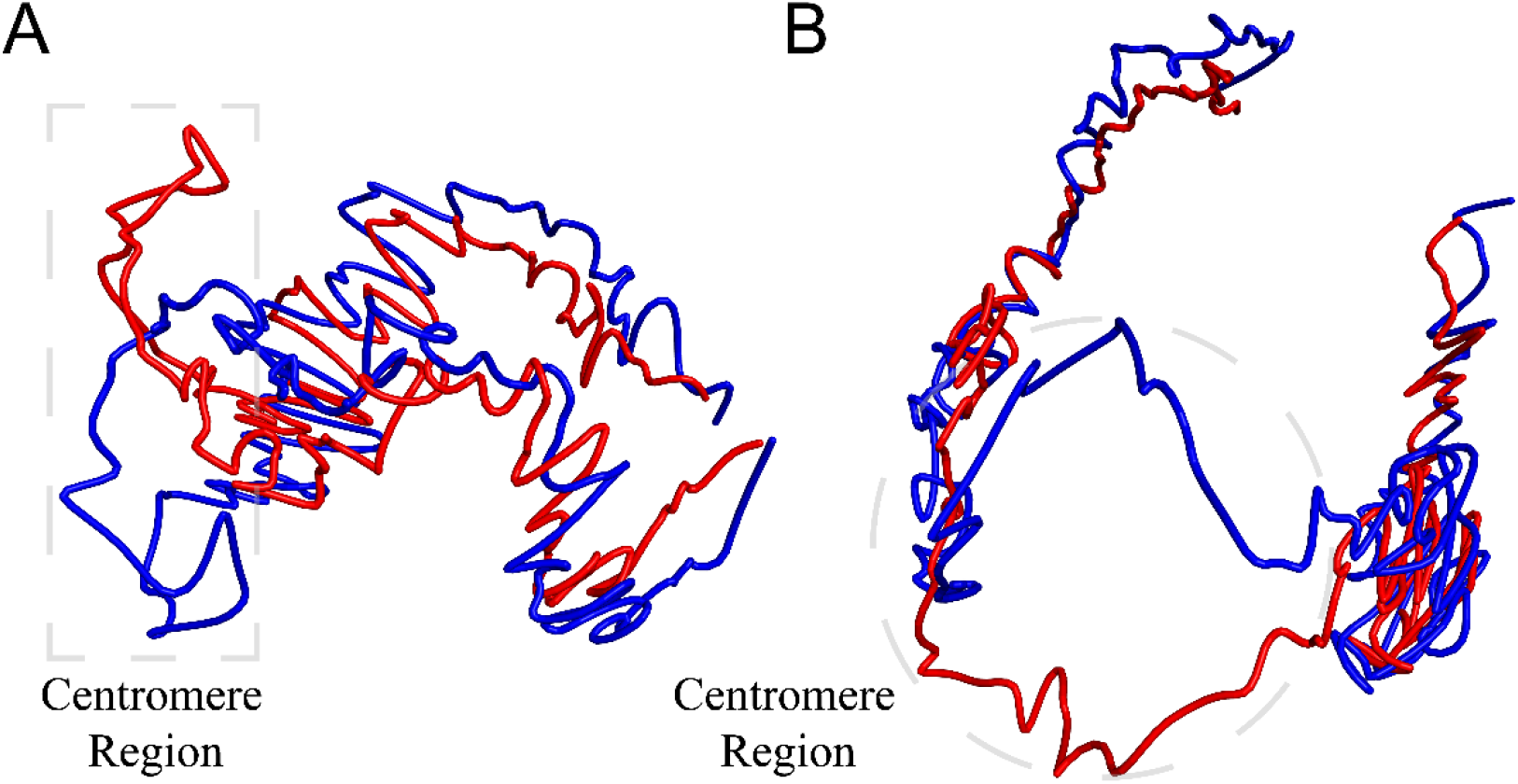
Comparison of the chromosome 1 structure between wild-type and AtMORC6 mutant of *Arabidopsis thaliana*. The red structure represents the wild-type chromosome and the blue structure represents the AtMORC6 mutant chromosome. A clear separation in the centromere region can be observed. (A) and (B) show the same structure in different orientations (viewpoints).

## 4. Discussion

In this paper, we have presented an algorithm called EVRC, which can be used to reconstruct the 3D structure of chromosomes based on spatial distance constraints that are related to chromatin condensation characteristics. The algorithm takes the co-clustering coefficient between two bins as the weight and sums the error vectors to determine the optimal coordinates of bin position. Through continuous iterative optimization, the error vector becomes smaller and smaller, and the reconstructed structure approaches the actual structure. We applied the EVRC algorithm to six simulating structure datasets of varying complexity and to human Hi-C experimental datasets. The reconstruction process of the algorithm was demonstrated, and its performance was evaluated using the Root Mean Square Deviation (RMSD) and Pearson Correlation Coefficient (PCC).

We also applied the EVRC algorithm to the Hi-C datasets of *Arabidopsis Thaliana*, and the reconstruction results of the five *Arabidopsis* chromosomes at different resolutions showed that the overall morphology of chromosomes was consistent. At low resolution, the reconstructed structure is smoother and displays the rough conformation of chromosomes, whereas at high resolution, the reconstructed structure shows more details. This indicates that the EVRC algorithm can accurately reconstruct chromosome conformation at different resolutions, and the algorithm is stable. We also reconstructed chromosome No. 1 of wild-type and mutant *Arabidopsis thaliana*, and found that the centromere regions are separated by 180°. In the mutant, the centromere region was closer to the surrounding chromatin, and the interaction was enhanced, resulting in the whole chromosome exhibiting a “C” shape. In contrast, in the wild-type, the centromere region folds upward, resulting in the chromosome exhibiting an “S” shape. These results confirmed the effect of AtMORC6 protein on the centromere region of chromosomes.

In summary, our EVRC algorithm has wide applicability and high accuracy, and has a great potential for multi-chromosome 3D structure reconstruction.

## Code Availability

https://github.com/mbglab/EVRC.

## Acknowledgements

This work was supported by the National Natural Science Foundation of China (Grant 31971184). The funders had no role in study design, data collection and interpretation, or the decision to submit the work for publication. Help from Kang-Jian Hua is highly appreciated.

## Supplementary materials

**Figure S1.**
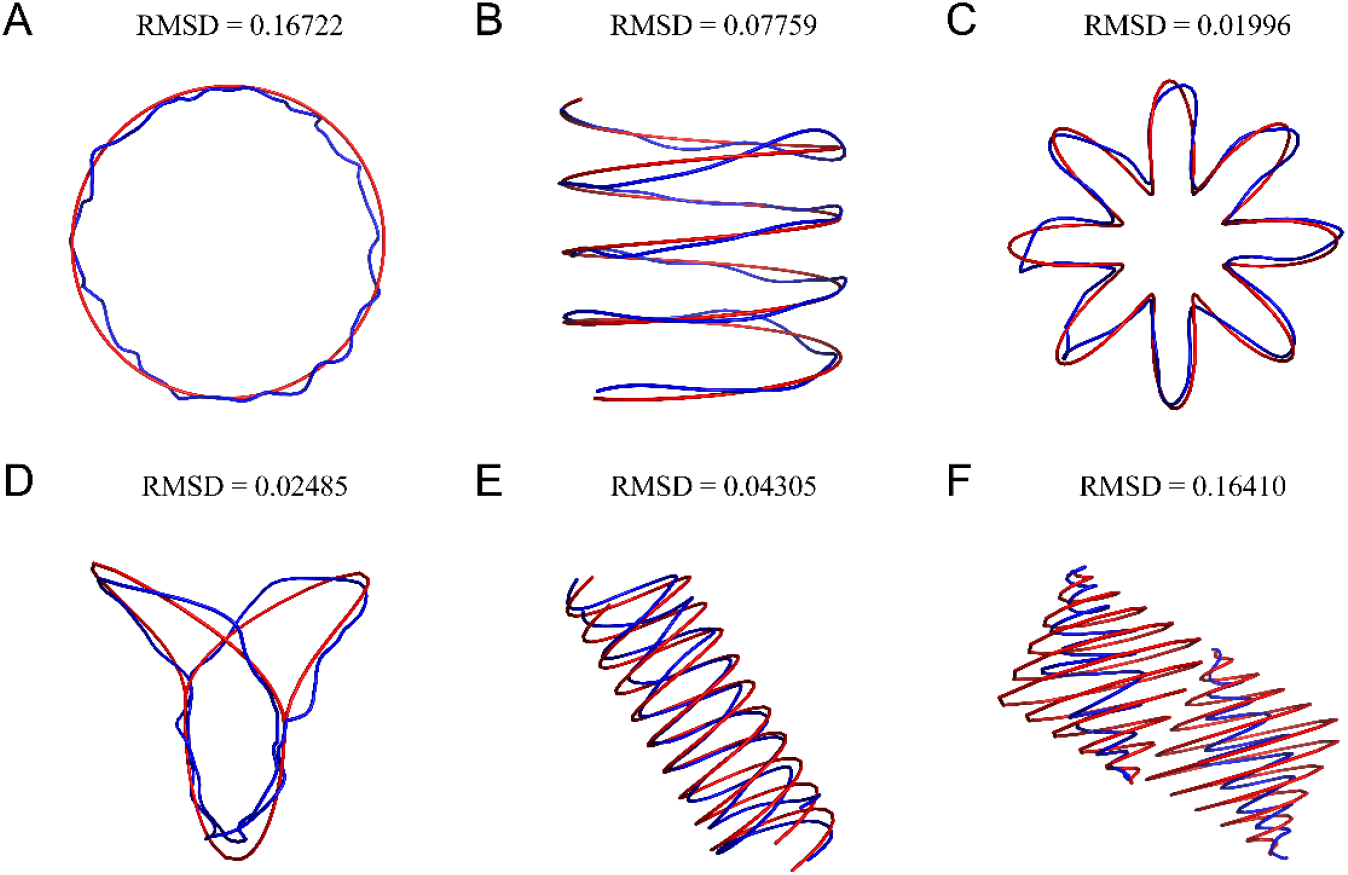
Reconstructed structure (blue curve) and original structure (red curve) when noise level is 1 and smoothing factor is 2. (A) circular curve (circle); (B) open spiral curve (spiral); (C) closed spiral curve (circular spiral); (D) replication fork; (E) double helix; (F) double spherical helix.

